# An EEG-based Multi-Scale Hybrid Attention and Squeeze Network for objective taste assessment

**DOI:** 10.1101/2025.11.06.686898

**Authors:** Tianyi Yang, Yueyue Xiao, Zhaoyan Li, Chunxiao Chen, Fei Ye, Mian Cao, Rui Liu, Yingli Zhu, Zhiyu Qian, Jagath C. Rajapakse

## Abstract

Taste perception is central to flavor experiences. Electroencephalography (EEG) signals carry rich information about taste perception. Because these neural data are independent of verbal reports and bypassing conscious filtering, EEG holds promise as a neural-sensing platform for objective taste representation. However, taste-evoked EEG signals exhibit complex spatiotemporal and spectral dynamics that require advanced computational approaches to decode effectively. This study developed an EEG-based Multi-Scale Hybrid Attention and Squeeze Network (MHASNet) for taste evaluation. EEG signals evoked by distinct taste stimuli were recorded, a dedicated taste-EEG dataset was compiled, and a novel deep learning architecture, MHASNet, was designed to classify these signals. MHASNet synergistically integrates multi-scale convolutions to capture temporal dynamics across different time scales, dual-attention mechanisms to localize discriminative brain regions and electrode positions, and a squeeze-and-excitation module to optimize frequency-band contributions-collectively enabling precise extraction of taste-specific neural signatures. Results showed that the proposed model achieved superior performance across five taste categories (sweet, sour, salty, bitter, and tasteless), with 94.33% accuracy, 91.37% F1-score, 91.89% precision, and 92.07% recall. These results surpass benchmark models while maintaining millisecond-level inference latency suitable for real-time applications. By complementing subjective evaluations and instrumental analyses, the model offers an objective, neurophysiology-based solution for taste evaluation.

## 1. Introduction

Taste is one of the primary sensory modalities through which humans interact with their external environment and serves as a critical component of flavor perception in food experiences (Shepherd, 2011). Sour, sweet, bitter, and salty are the most universally recognized basic taste qualities, with complex taste experiences often interpreted as combinations of these elements (Bartoshuk, 1978; Beauchamp, 2019). Taste perception is initiated when chemical compounds interact with specific chemoreceptors on the tongue, activating receptor cells within taste buds (Chandrashekar et al., 2006). These chemosensory signals are conveyed via cranial nerves, such as the facial and glossopharyngeal nerves, to the nucleus of the solitary tract. From there, they are relayed to the thalamus and then projected to the primary gustatory cortex (Pritchard et al., 1986; Rolls, 2020). Within the central nervous system, distinct taste stimuli evoke specific activation patterns in cortical regions such as the insula and orbitofrontal cortex, contributing to the diversity and subjectivity of taste-related perceptual experiences (Shepherd, 2011; Yang et al., 2023). However, these electrophysiological responses are rapid and multidimensional, involving hierarchical signal processing from peripheral sensory interfaces to central integration networks, which makes them difficult to characterize precisely using conventional sensory evaluation approaches or artificial sensing technologies such as electronic tongues.

Exploring brain activity provides new insights into sensory experience. Since the brain is the core organ responsible for perception and decision-making, neural recordings can more authentically reflect true consumer experiences. Such objective neural data are independent of verbal reports. They capture responses to tasting, smelling, or viewing food and bypass conscious filtering (Songsamoe et al., 2019). Our previous electroencephalography (EEG) studies show distinct electrophysiological signatures and perceptual characteristics for different taste stimuli. Each stimulus shows unique temporal dynamics and signal response profiles in the brain, especially in gustatory regions such as the insula and the orbitofrontal cortex (Yang et al., 2023, 2024). These neural signals carry rich information about sensory coding and subjective experience. Therefore, EEG has the potential to serve as a noninvasive neural-sensing platform that objectively represents taste perception, with broad application prospects.

For instance, EEG-driven models objectively assess flavor profiles. They reduce reliance on subjective panels and minimize evaluation variability (Songsamoe et al., 2019). Moreover, measuring and analyzing neural responses to different tastes can enhance consumer research. For example, neural patterns of olfactory-induced sweetness enhancement can guide flavor combinations that amplify perceived sweetness without added sugar (Xia et al., 2025). This data-driven strategy for precision product development improves design accuracy. It allows developers to tailor and optimize flavor attributes using quantitative neural feedback rather than trial-and-error. For individuals with gustatory impairment, the same technologies inform assistive devices that substitute or augment taste perception (Crofton et al., 2019). At the health interface, analysis of EEG bio-signals may facilitate the early screening of disorders commonly linked to gustatory dysfunction, including cancer, diabetes, anorexia, and COVID-19 (Cui et al., 2023).

However, accurately extracting taste-related information from the noisy and complex nature of EEG signals remains a significant challenge. EEG signals exhibit low signal-to-noise ratios and are vulnerable to artifacts from facial muscle movements, eye blinks, and head motion during tasting (Urigüen & Garcia-Zapirain, 2015). Taste-evoked responses show high inter-individual variability and are distributed across multiple frequency bands (δ-γ) and brain regions without unified extraction standards (Yang et al., 2023). Additionally, handcrafted features struggle to capture the nonlinear, multidimensional neural dynamics underlying taste perception, where transient event-related potentials coexist with sustained oscillatory activities across different temporal scales (Makeig et al., 2004; Cui et al., 2023).

In recent years, rapid advancements in deep learning have resulted in transformative progress in EEG signal analysis. Unlike traditional methods that rely on handcrafted features, deep learning models can automatically learn high-level, nonlinear, and discriminative representations directly from raw EEG data, enabling more effective modeling of its complex spatiotemporal dynamics (Hossain et al., 2023). For example, convolutional neural networks leverage local connectivity and weight sharing to efficiently extract localized features and integrate global context through multiple convolutional layers, demonstrating strong spatiotemporal modeling capabilities (Islam et al., 2022). Transformer-based models, by incorporating self-attention mechanisms, capture long-range dependencies across both time and space without fixed receptive fields. This makes them well-suited for identifying non-local neural patterns and salient spatiotemporal features in EEG signals (Khan et al., 2022).

Despite these advantages, most studies have focused primarily on domains such as emotion recognition and motor imagery (Fernandes et al., 2024), with limited attention given to taste perception based on EEG decoding. Recent efforts include channel-reduction approaches that achieve 97.85% accuracy with 12 selected channels but prioritize computational efficiency over spatial completeness (Xia et al., 2025), and conventional wavelet-based feature extraction methods that reach 83.62% accuracy for binary classification but lack hierarchical modeling capacity (Vo et al., 2023). These approaches provide limited mechanistic insights into the underlying neurophysiological basis of taste perception. They either sacrifice spatial resolution or fail to simultaneously address the temporal, spatial, and spectral complexity of taste-evoked neural responses. Taste-related EEG involves complex interactions across frequency, time, and spatial dimensions (Yang, et al., 2023, 2024). General-purpose models often struggle to capture unique neural representations and remain limited in multi-scale feature extraction and semantic integration.

To address these challenges, we developed an EEG-based Multi-Scale Hybrid Attention and Squeeze Network (MHASNet) for taste evaluation. MHASNet integrates parallel multi-scale temporal convolutions to capture both transient and oscillatory EEG patterns, hierarchical attention mechanisms (channel and spatial) to localize taste-relevant cortical regions and electrodes, and a squeeze-and-excitation (SE) module to reweight δ–γ frequency bands for enhanced spectral discriminability. This study demonstrates the potential of integrating EEG signals with deep-learning methods for taste evaluation. By precisely capturing EEG features related to taste perception, it enables objective, real-time quantification of taste experience.

## 2. Materials and methods

### 2.1 Participants

Twenty-five volunteers participated in this study after providing written informed consent. The inclusion criteria were as follows: all participants were right-handed, were not taking medications, had no history of psychiatric or neurological disorders, and had no professional background in taste discrimination. Five participants were excluded from the analysis: three due to failure to correctly identify more than 66% of the taste stimuli or reporting weak taste perception (intensity rated below 50 on a 100-point scale), and two due to unusable EEG recordings. As a result, 20 participants (12 males and 8 females; mean age = 23 years, SD = 2.6 years) were included in the final analysis. The study protocol was approved by the Research Ethics and Technical Safety Committee of Yangzhou University (Reference number: YXYLL-2024-113), and was conducted in accordance with the ethical principles outlined in the latest revision of the Declaration of Helsinki.

### 2.2 Taste stimuli

The selection of gustatory stimuli was based on our previously established protocol for taste research (Yang et al., 2023). All compounds used have been identified as characteristic representatives of the four basic taste modalities and were confirmed to be reliably distinguishable by participants. The specific concentrations were as follows: sucrose (1.0 M) as the sweet stimulus; quinine (1.0×10⁻³ M) as the bitter stimulus; citric acid (0.1 M) as the sour stimulus; and sodium chloride (0.5 M) as the salty stimulus. A tasteless artificial saliva solution (pH 6.8) was obtained from a local retail pharmacy. All substances were of food-grade quality.

### 2.3 EEG acquisition system

EEG data were acquired with the NeuroScan NuAmps Express system (32 channels), and real-time signal visualization and impedance monitoring were conducted using Curry 7 software (NeuroScan, USA). Electrodes were positioned according to the international 10–20 system, with FCz designated as the reference electrode to reduce global scalp-potential shifts and enhance signal stability (Yang et al., 2023). The sampling rate was fixed at 1000 Hz. During acquisition, signal quality and electrode impedances were continuously monitored in real-time through the Curry 7 impedance check interface and waveform display. When impedance exceeded 5 kΩ or significant artifacts were detected, the recording was paused, and the experimenter manually re-prepared the affected electrodes by adding conductive gel or adjusting electrode placement before resuming data collection. Throughout the experiment, electrode impedance was maintained below 5 kΩ to ensure reliable signal transmission and minimize artifacts.

Fig. 1 depicts the complete workflow of the taste-EEG evaluation. Following the presentation of each taste stimulus, EEG signals from multiple cortical regions are acquired via an electrode cap, transmitted through shielded cables to a signal-amplifier module, and simultaneously rendered for visual inspection. The signals undergo standardized preprocessing and are subsequently fed into the custom deep learning architecture, MHASNet, for classification and identification. The digital electrode layout employed in this study is presented in Fig. 2. A comprehensive description of the EEG acquisition and recording procedures is supplied in the *Supplementary Information* (Fig. S1).

**Fig. 1.**
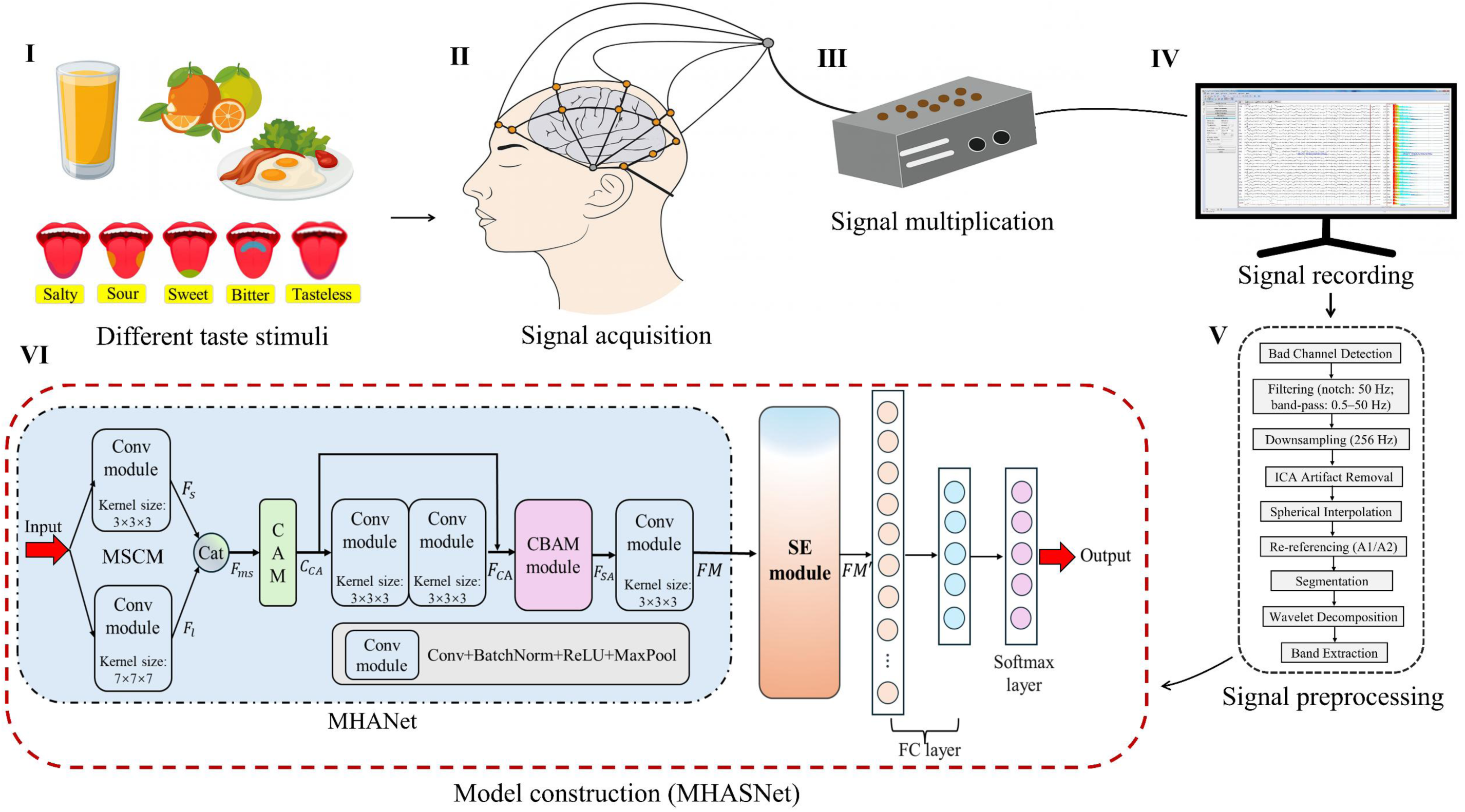
Schematic workflow of the MHASNet-based taste-EEG evaluation.

**Fig. 2.**
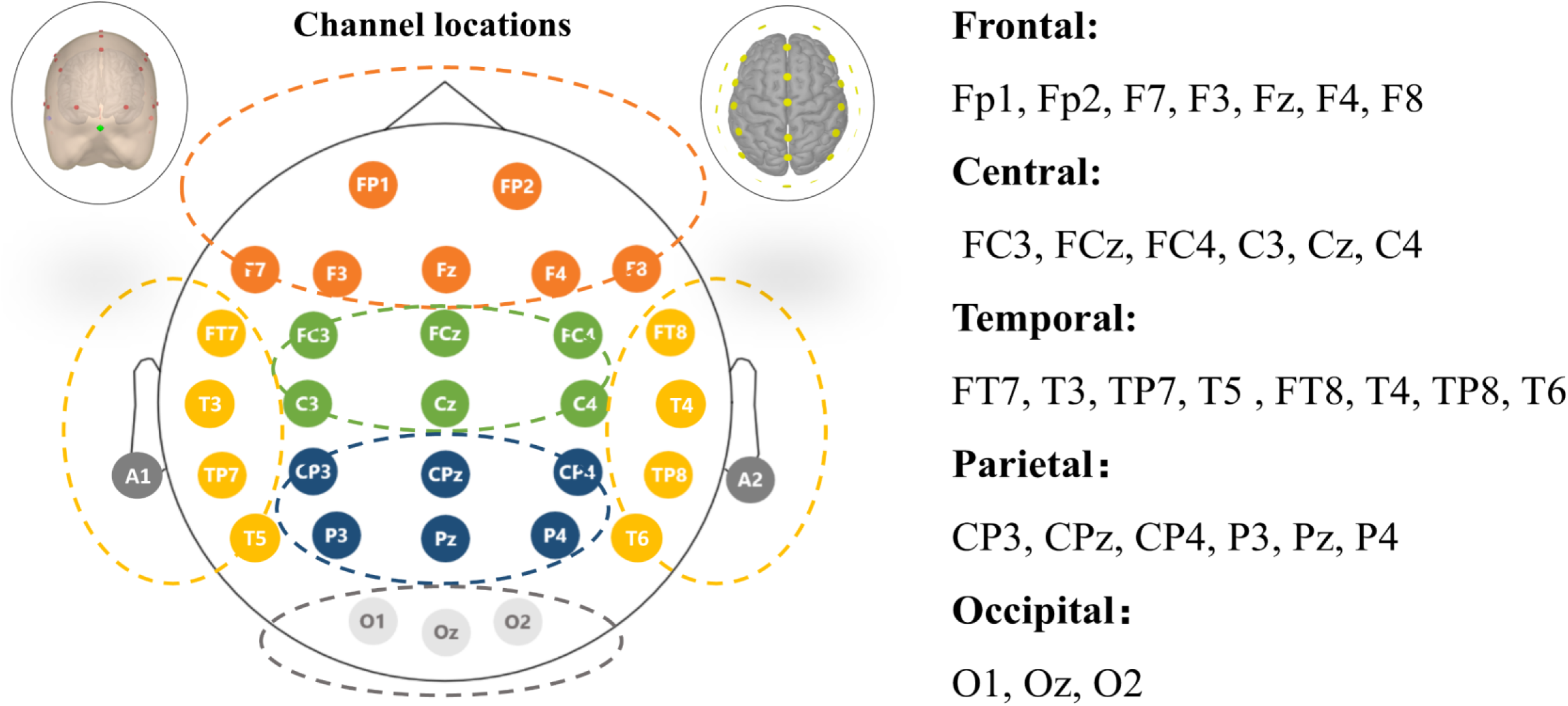
The EEG cap layout for 32 electrodes. Electrodes are arranged according to the international 10–20 system and are grouped into five anatomical regions: frontal (orange), central (green), temporal (yellow), parietal (blue), and occipital (gray). Reference electrodes A1 and A2 were placed on the left and right mastoids, respectively.

### 2.4 Data preprocessing

Offline EEG data were preprocessed using MNE-Python. Before filtering, bad channels were identified through automatic detection (peak-to-peak amplitude >±150 μV in raw data) followed by visual confirmation. Channels exhibiting persistent artifacts, flat signals, or excessive noise were excluded. A notch filter (50 Hz, bandwidth: 2 Hz) was then applied to remove power-line interference, followed by a finite-impulse-response band-pass filter (0.5–50 Hz) to suppress low-frequency drifts and high-frequency noise. The filtered data were then downsampled to 256 Hz. Independent component analysis (ICA) was used to remove ocular, muscular, and cardiac artifacts. To preserve component independence, bad channels were excluded before ICA decomposition. After artifact correction, spherical-spline interpolation reconstructed the excluded channels, restoring the full electrode montage. Signals were re-referenced to the average of the left and right mastoids (A1 and A2) to reduce voltage offsets and enhance spatial uniformity.

After preprocessing, a sliding-window augmentation strategy was applied to enlarge the training dataset. Each taste-stimulus trial, which lasted 5 s (solution-holding phase), was segmented into nine overlapping 1-s windows with a 0.5-s overlap. To prevent data leakage, all segments from a given trial were allocated exclusively to either the training or the test set. Twenty participants each received five taste stimuli (sour, sweet, bitter, salty, and tasteless), repeated five times, yielding 25 trials per participant. With nine segments per trial, a total of 4,500 segments were generated (20×5×5×9). Each segment underwent six-level wavelet-packet decomposition using Daubechies wavelets, producing 64 sub-band components at approximately 2 Hz per sub-band (256 Hz sampling rate). Sub-bands corresponding to the physiologically relevant EEG frequency range (0.5–50 Hz)—approximately 25 sub-bands—were retained. These sub-bands were grouped according to five canonical frequency bands: δ (0.5–4 Hz), θ (4–8 Hz), α (8–13 Hz), β (13–30 Hz), and γ (30–50 Hz), and reconstructed through inverse wavelet packet transformation (Xia et al., 2023). For each segment, the multi-band data were reshaped into a tensor of size 5 × 30 × 256, where 5 represents frequency bands, 30 represents the active electrodes (excluding reference electrodes A1 and A2), and 256 represents the time points. These tensors were fed into the proposed MHASNet to classify the five taste categories.

### 2.5 Construction of the taste EEG classification model

An illustrative example of sour-evoked EEG signals is shown in Fig. 3, with left panels displaying multi-channel spatial distributions and right panels showing representative waveforms from electrode Fz for each frequency band. As illustrated, taste-evoked EEG signals exhibit complex and heterogeneous characteristics across temporal, spatial, and spectral domains. Consequently, the effective capture of these multidimensional neural dynamics is essential for accurate decoding and interpretation. To achieve this objective, we propose MHASNet, as depicted in the model-construction panel of Fig. 1. The architecture comprises two primary components: a Multi-scale Hybrid Attention sub-network (MHANet) and a SE module. Within MHANet, two key sub-modules are integrated: the Multi-Scale Convolutional Module (MSCM) and the Convolutional Block Attention Module (CBAM). Together, these modules enhance the decoding of taste-evoked EEG signals by adaptively capturing multi-scale contextual information and emphasizing salient neural features through attention mechanisms.

**Fig. 3.**
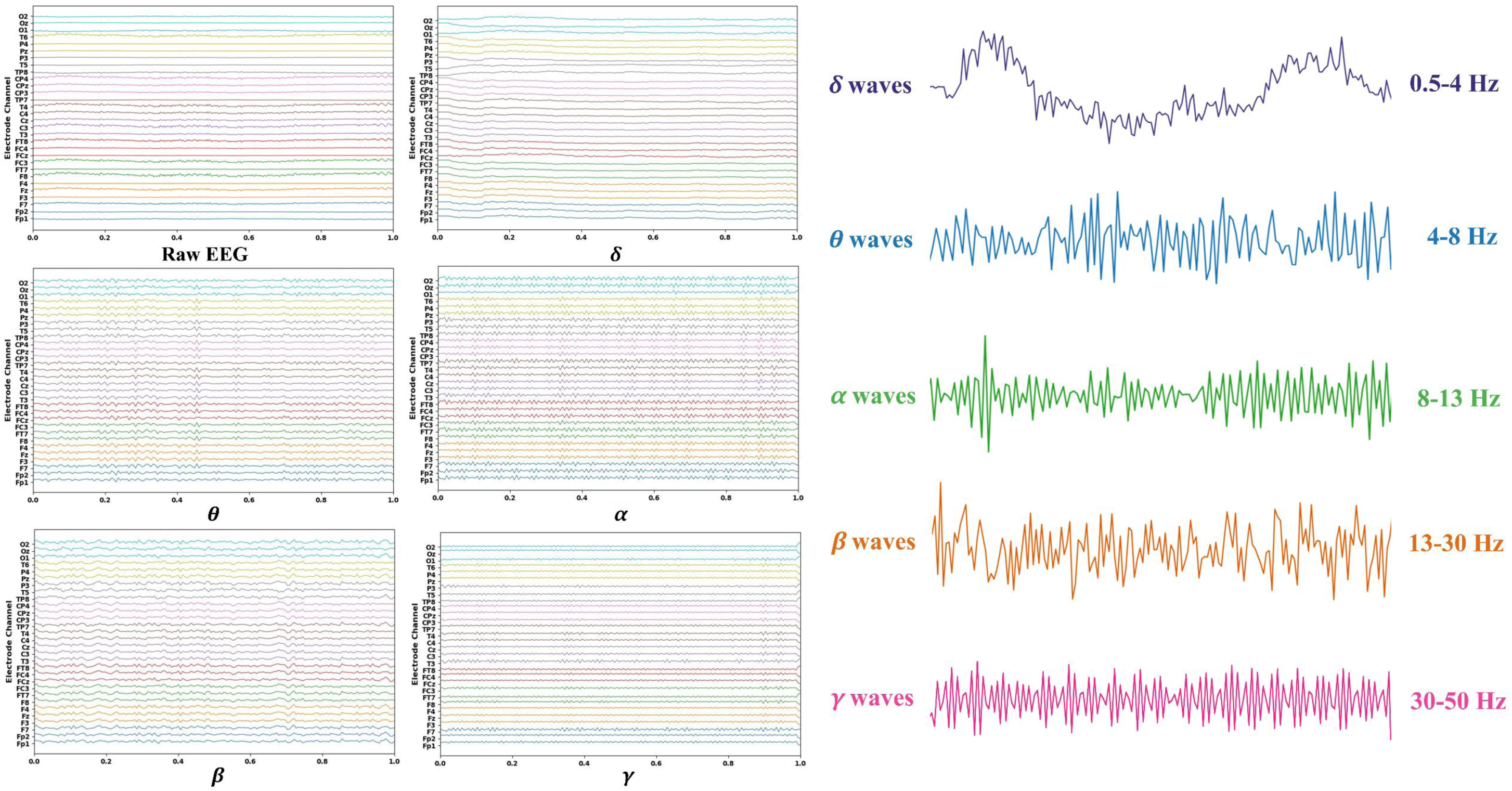
Sour-evoked gustatory EEG sample illustrating spatiotemporal and spectral heterogeneity. The top-left panel shows the raw EEG signal across 30 scalp channels over a 1-second window. The remaining left panels display the corresponding frequency-decomposed EEG signals using wavelet packet reconstruction for five canonical bands (δ, θ, α, β, and γ). The right column presents representative single-channel waveforms from electrode Fz for each frequency band, illustrating the characteristic oscillatory patterns.

#### 2.5.1 Parameter settings

All experiments were implemented in PyTorch (Python 3.9) and run on a workstation with an Intel Core i9-14900 CPU (3.59 GHz), 32 GB RAM, and an NVIDIA RTX 3060 (12 GB VRAM). We used the Adam optimizer with an initial learning rate of 0.001, a batch size of 10, and up to 200 epochs. These hyperparameters were selected based on common practices in EEG-based deep learning and preliminary validation on our dataset. Early stopping was used to limit overfitting by halting training if the validation loss did not improve for 15 consecutive epochs. The checkpoint with the best validation performance was retained.

#### 2.5.2 Multi-scale feature extraction

Inspired by neuroscience models that use multi-scale receptive fields to decode complex sensory inputs, we adopted a multi-scale convolutional strategy to capture short- and long-range temporal dependencies in EEG (Mohammadi Foumani et al., 2024). We applied two parallel three-dimensional convolutional kernels, sized 3×3×3 and 7×7×7, with receptive fields aligned to the temporal axis. The 3×3×3 kernel targets local fluctuations and transient components, such as rapid spikes or event-related potentials. The 7×7×7 kernel captures longer-lasting patterns and oscillatory trends across wider time windows. This design builds multi-resolution temporal features. It increases sensitivity to diverse neural response patterns evoked by taste stimuli.

Specifically, given an EEG input signal *X* ∈ ℝ^*a*×*b*×*c*^, where *a* = 5 is the number of frequency bands, *b* = 30 is the number of electrode channels, and *c* = 256 is the time step, the multiscale convolution is computed as follows:

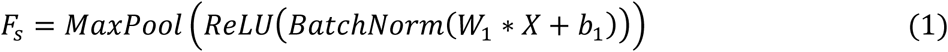

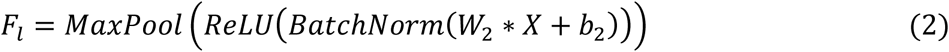

where W_1_ and W_2_ denote the weights of the 3×3×3 and 7×7×7 convolution kernels, respectively, b_1_ and b_2_ are bias terms. The features obtained from short-term convolution (3×3×3) and long-term convolution (7×7×7) are concatenated, i.e., F_ms_ = Cat(F_s_, F_l_), to preserve both local and global information, thereby enhancing the model’s ability to capture temporal dependencies.

#### 2.5.3 Attention module

To capture the spatial heterogeneity of taste-evoked EEG, MHASNet uses a two-stage attention refinement across channel and spatial dimensions. This design encodes both global anatomical relevance and localized spatial saliency in multichannel data (Woo et al., 2018). In the first stage, a Channel Attention Module (CAM) is applied. CAM adaptively recalibrates feature weights across EEG channels according to task-specific relevance. Because each channel corresponds to a distinct cortical region, CAM emphasizes informative responses from areas more sensitive to gustatory stimulation. As illustrated in Fig. 4a, global average pooling and max pooling are applied along the temporal axis to summarize the activation patterns of each channel. The resulting descriptors are passed through a shared multilayer perceptron (MLP), followed by a sigmoid activation, to generate channel-wise attention weights:

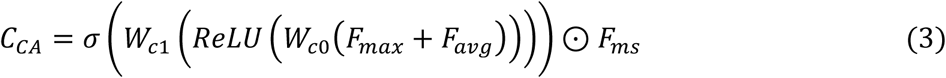

**Fig. 4.**
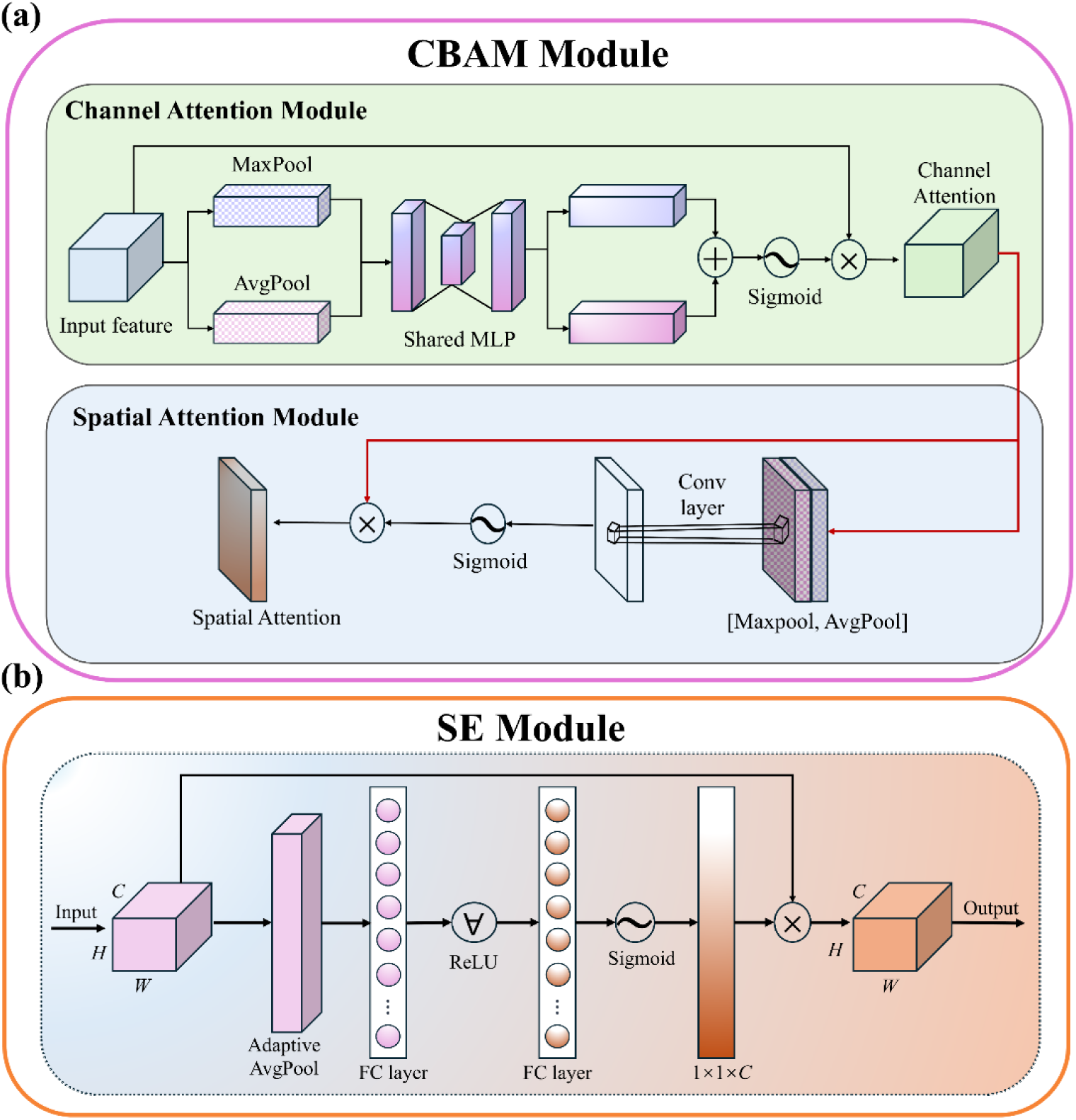
Architectures of (a) the CBAM and (b) the SE module.

where *W*_*c*0_ and *W*_*c*1_ are the weights of the MLP, ⊙ represents element-wise multiplication, *F*_*max*_ = *MaxPool*(*F*_*ms*_), *F*_*avg*_ = *AvgPool*(*F*_*ms*_), and *σ* denotes the sigmoid activation function.

While CAM offers a coarse-level prioritization across cortical regions, it lacks the ability to capture fine-grained spatial variation across the electrode topology. To address this limitation, MHASNet integrates a CBAM, which introduces an embedded Spatial Attention Module (SAM) to extend attention refinement to the spatial dimension. SAM first aggregates spatial features by averaging and max pooling along the channel axis, followed by a convolutional layer and sigmoid activation to produce a spatial attention map. This map enables the model to dynamically highlight salient electrode locations that exhibit discriminative activation patterns in response to specific taste stimuli, as expressed by:

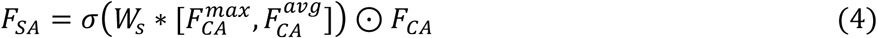

where *F*_*CA*_ = *CM*(*CM*(*C*_*CA*_)), *CM* represents the Conv module, *W*_*s*_is the convolution kernel weight, ∗ represents the convolution operation. *σ* is the *sigmoid* activation function, which ensures that recalibration weights remain within (0,1). 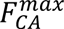 and 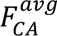 denote the maximum pooling and average pooling results on the channel dimension, respectively, 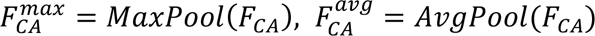.

By sequentially integrating CAM and CBAM, MHASNet establishes a hierarchical spatial attention structure: CAM determines which cortical regions are relevant, while SAM further identifies where within those regions the most salient neural activity occurs. This joint refinement enhances the model’s capacity to capture the spatial characteristics of taste-evoked EEG responses.

#### 2.5.4 Frequency-specific feature recalibration via SE module

To address the challenge of suboptimal frequency-band feature selection in EEG analysis (Yang et al., 2023), MHASNet incorporates a SE module (Fig. 4b). The SE module uses global contextual information to adaptively recalibrate the importance of each frequency-band channel. This refinement enhances extraction of discriminative representations from multiband EEG inputs (Woo et al., 2018).

The module operates in two stages: squeeze and excitation. In the squeeze phase, the spatial and temporal dimensions of the input feature map *FM* ∈ ℝ^*H*×*W*×*C*^ are aggregated using global average pooling, where *H* and *W* denote the height and width of the feature map, and *C* is the number of channels. This operation yields a compact descriptor for each channel:

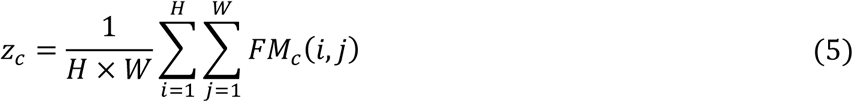

where *z*_*c*_ represents the statistical summary of the *c*-th channel. This global descriptor captures holistic activation information and serves as the basis for adaptive weighting.

In the excitation phase, the aggregated vector is passed through a bottleneck structure consisting of two fully connected layers. The dimensionality is first reduced to *C*/*τ*, where *τ* is a reduction ratio, followed by a ReLU activation and a sigmoid function. This structure captures channel-wise dependencies and generates refined attention weights:

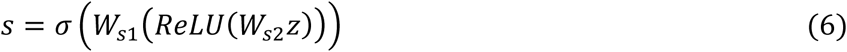

where *W*_*s*1_ ∈ ℝ^*C*/*τ*×*C*^ and *W*_*s*2_ ∈ ℝ^*C*×*C*/*τ*^ are trainable weight matrices. This excitation mechanism models inter-channel dependencies and generates refined weights for frequency-specific recalibration.

The final scaling operation applies the learned attention weights *s* to the original feature map through element-wise multiplication:

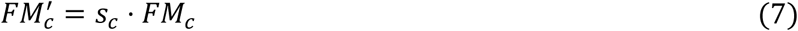

where 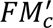 represents the recalibrated feature map for the *c*-th channel. This frequency-aware weighting mechanism adaptively enhances the contribution of the most informative components while suppressing noisy or less relevant channels. By integrating the SE module, MHASNet improves its sensitivity to task-relevant spectral patterns.

#### 2.5.5 Loss functions

In the MHASNet framework, the cross-entropy loss is employed as the primary optimization objective to guide the learning process. As a widely used loss function for multi-class classification tasks, cross-entropy effectively quantifies the discrepancy between the predicted class probabilities and the true labels, enabling the network to iteratively adjust its parameters to improve classification performance (Connor et al., 2024). The mathematical formulation of the cross-entropy loss is defined as:

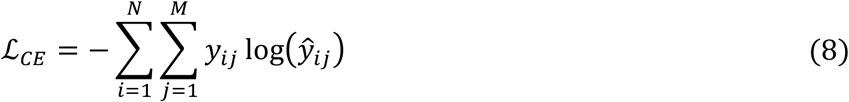

where *N* is the number of samples, *M* denotes the total number of classes, *y*_*ij*_ is the ground truth label for class *j* of sample *i* (one-hot encoded), *ŷ_ij_* represents the predicted probability of sample *i* belonging to class *j*, computed using the softmax function:

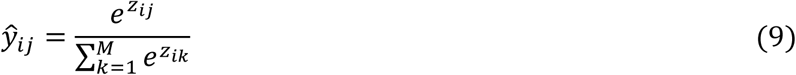

where *z*_*ij*_ denotes the raw model output for class *j*.

During training, the cross-entropy loss drives the model to maximize the predicted probability of the correct class while minimizing those of incorrect classes, thereby enabling MHASNet to extract discriminative features from taste-evoked EEG signals and improve classification performance in taste perception decoding.

#### 2.5.6 Performance evaluation

To comprehensively evaluate the classification performance of the model, four standard metrics were employed: Accuracy, Precision, Recall, and F1-score. An 80:20 train–test split with five-fold cross-validation was implemented to ensure robust evaluation. In each fold, 80% of the data was used for training and 20% for validation, ensuring that model performance was assessed on previously unseen samples. All reported results are averaged over the five folds.

## 3. Results and Discussion

### 3.1 Comparison with state-of-the-art methods

We validate MHASNet by comparing it with seven state-of-the-art EEG classifiers: EEGNet (Lawhern et al., 2018), ResNet18 (Bagherzadeh et al., 2024), DenseNet (Pusarla et al., 2022), EfficientNetV2 (Tan & Le, 2021), MobileNetV2 (Sandler et al., 2018), FBANet (Xia et al., 2023), and LGFNet (Xia et al., 2025). The evaluation covers four metrics: accuracy, F1-score, precision, and recall—and also inference time as a measure of efficiency.

Table 1 summarizes the quantitative results. MHASNet attains the highest accuracy at 94.33%, significantly outperforming all baseline models. It also shows superior F1-score (91.37%), precision (91.89%), and recall (92.07%), indicating robust and reliable decoding of taste-evoked EEG. Among competitor models, FBANet reaches 91.22% accuracy but still lags behind across all metrics. LGFNet (88.11%), MobileNetV2 (87.78%), and EfficientNetV2 (87.56%) perform competitively yet below MHASNet. ResNet18 and DenseNet perform moderately, and EEGNet records the lowest accuracy. In addition to classification effectiveness, inference time is also critical for real-time decoding. MHASNet achieves 2.31 ms, which is comparable to MobileNetV2 (2.21 ms) and notably faster than FBANet (5.31 ms), LGFNet (5.03 ms), and DenseNet (4.31 ms). These results support the potential of MHASNet for efficient, real-time applications in taste perception decoding.

**Table 1.** Quantitative comparison of the performance of our method against other state-of-the-art methods.

| Method | Accuracy (%) | F1-score (%) | Precision (%) | Recall (%) | Time (ms) |
| --- | --- | --- | --- | --- | --- |
| EEGNet | 81.44±0.16 | 73.88±0.23 | 75.12±0.23 | 75.89±0.23 | 0.76 |
| ResNet18 | 84.89±0.16 | 77.57±0.23 | 78.74±0.23 | 79.82±0.23 | 6.31 |
| DenseNet | 85.56±0.15 | 79.18±0.22 | 80.70±0.21 | 80.20±0.22 | 4.31 |
| EfficientNetV2 | 87.56±0.15 | 82.59±0.20 | 84.21±0.20 | 83.41±0.20 | 3.18 |
| MobileNetV2 | 87.78±0.14 | 82.74±0.21 | 83.84±0.21 | 84.22±0.20 | 2.21 |
| FBANet | 91.22±0.13 | 87.07±0.20 | 87.66±0.19 | 88.03±0.19 | 5.31 |
| LGFNet | 88.11±0.15 | 84.13±0.20 | 85.40±0.19 | 85.11±0.20 | 5.03 |
| MHASNet | 94.33±0.10 | 91.37±0.16 | 91.89±0.15 | 92.07±0.15 | 2.31 |

Confusion matrices in Fig. S2 further illustrate that MHASNet achieves higher true positive rates and lower misclassification rates across all taste categories compared to other models, reinforcing its ability to capture discriminative neural signatures. These results show that MHASNet achieves an effective trade-off among classification accuracy, computational efficiency, and robustness. They also show that it outperforms conventional deep networks and domain-specific architectures such as FBANet and LGFNet. The model’s consistent performance across multiple metrics validates its potential for practical deployment in EEG-based taste evaluation platforms.

### 3.2 Ablation experiment of MHASNet

We conducted a progressive ablation study to assess how each MHASNet component contributed to performance in classifying taste-evoked EEG. Modules were integrated sequentially, and their individual and cumulative impacts on classification were evaluated. Fig. 5 presents a comparison of classification accuracy across model variants. The baseline, MHASNet-O, was a conventional CNN that excluded the multi-scale convolution, hybrid attention, and SE modules. It attained 87.22% accuracy and served as the reference configuration.

**Fig. 5.**
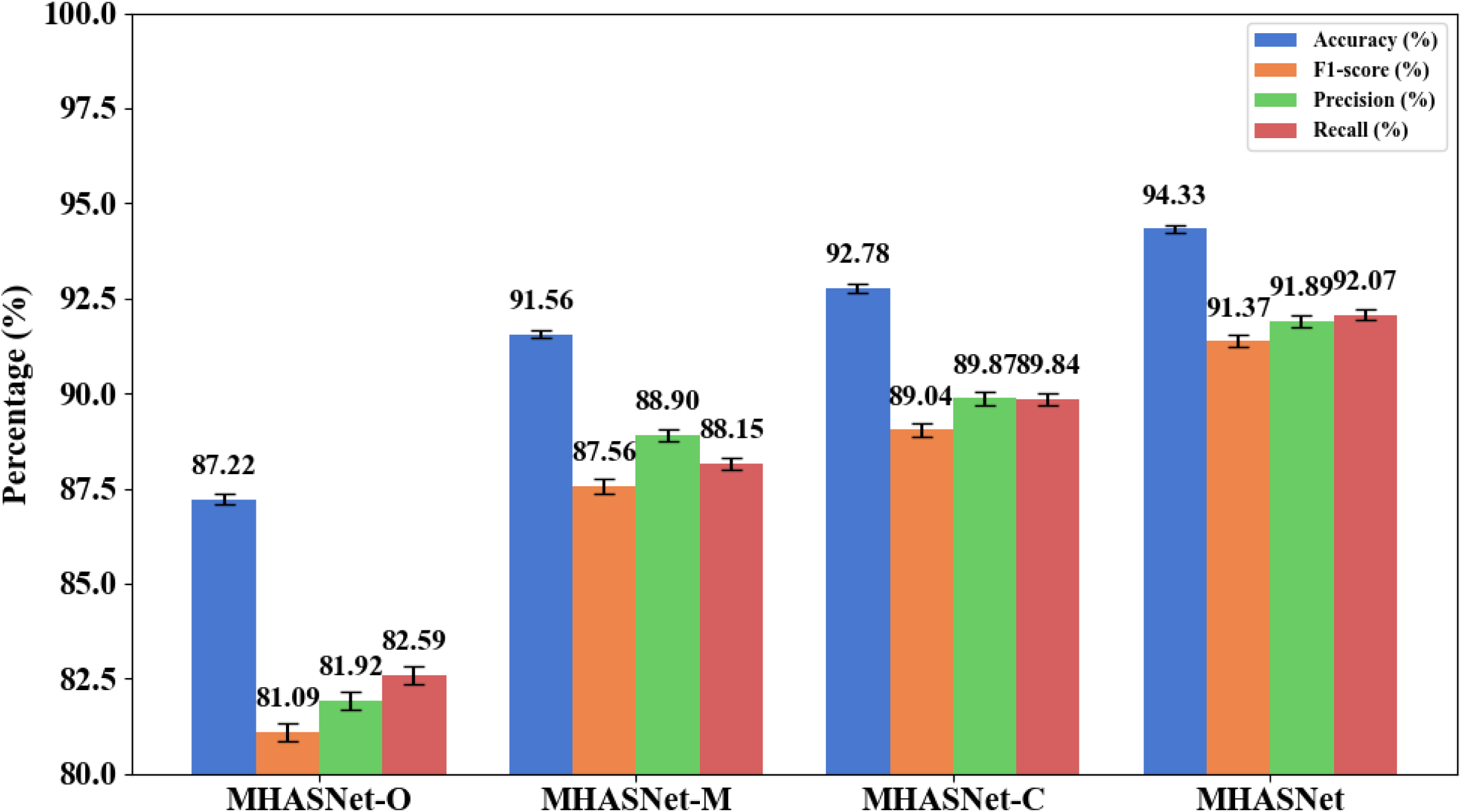
Results of ablation experiment. This figure presents the progressive performance improvements of MHASNet through stepwise integration of its architectural components. Performance is evaluated using four metrics: Accuracy, F1-score, Precision, and Recall.

Introducing multi-scale convolution and CAM (MHASNet-M) substantially improved performance, raising accuracy to 91.56%. This result confirmed the effectiveness of combining temporal multi-scale modeling with adaptive channel-wise weighting for capturing task-relevant EEG dynamics. Further incorporating CBAM yielded MHASNet-C and raised accuracy to 92.78%, thereby highlighting the value of spatial attention in emphasizing discriminative electrode regions and refining feature representations. Finally, integrating the SE module produced the complete MHASNet, which achieved 94.33% accuracy. By adaptively reweighting frequency-band components, the SE module enhanced robustness and spectral discriminability; improvements were also reflected in F1 = 91.37%, precision = 91.89%, and recall = 92.07%. These progressive gains validated the synergy among multi-scale feature extraction, hierarchical attention, and frequency-aware recalibration. Together, these components enhanced MHASNet’s ability to decode taste perception from EEG with high robustness and accuracy.

### 3.3 MHASNet-based validation and exploration of taste perception mechanisms

Previous studies show that taste perception elicits characteristic neural oscillations and region-specific cortical activity (Yang et al., 2023, 2024, 2025). In this section, we test whether the MHASNet-based taste-EEG model captures these established spectral and spatial hallmarks. The analysis showed strong agreement with known neurophysiological signatures. It also reveals how distinct taste qualities recruit specific oscillatory dynamics and cortical regions. These results provide mechanistic evidence for the model’s neurophysiological fidelity. They also extend understanding of the neural processes through which different tastes shape consumer preferences.

#### 3.3.1 Unveiling hierarchical spectral dynamics of taste perception via MHASNet

To further investigate and validate the contribution of distinct EEG frequency bands to taste-perception decoding, we performed a comparative analysis using features from individual bands—δ (0.5–4 Hz), θ (4–8 Hz), α (8–13 Hz), β (13–30 Hz), and γ (30–50 Hz)—and from their combined representation. Table 2 presents the classification metrics (accuracy, F1-score, precision, and recall) for each band and for the full-spectrum multi-band feature set.

**Table 2.** Taste classification results for single-frequency and multi-frequency EEG features.

| Method | Accuracy (%) | F1-score (%) | Precision (%) | Recall (%) |
| --- | --- | --- | --- | --- |
| $\delta$ | 92.44±0.11 | 89.56±0.16 | 90.40±0.15 | 90.59±0.15 |
| $\theta$ | 92.33±0.12 | 88.79±0.18 | 89.15±0.18 | 89.80±0.17 |
| $\alpha$ | 91.11±0.12 | 87.14±0.18 | 87.87±0.18 | 88.40±0.17 |
| $\beta$ | 90.56±0.13 | 85.81±0.19 | 86.55±0.19 | 86.79±0.18 |
| $\gamma$ | 90.89±0.13 | 86.59±0.20 | 87.37±0.19 | 87.64±0.19 |
| All | 94.33±0.10 | 91.37±0.16 | 91.89±0.15 | 92.07±0.15 |

Specifically, features derived from individual bands achieved reasonably good classification performance. Among the single-band models, the δ and θ bands yielded the highest accuracies, reaching 92.44% and 92.33%, respectively, with corresponding F1-scores exceeding 88%. These results suggest that low-frequency oscillations, particularly δ and θ rhythms, may carry critical discriminative information relevant to taste perception, potentially reflecting early-stage gustatory sensory encoding and attentional modulation processes (Cui et al., 2023; Wallroth et al., 2018).

The α band also demonstrated strong discriminative capacity, achieving an accuracy of 91.11% and an F1-score of 87.14%. Given that α-band activity is associated with cognitive modulation, sensory integration, and emotional processing, its contribution may reflect the integration of sensory identity with affective appraisal (Knyazev, 2007; Kotini et al., 2016). Considering the intrinsic coupling between taste perception and affective evaluation, where stimuli such as sweetness and bitterness evoke innate appetitive or aversive responses, α-band signals are likely to encode not only sensory identity but also the emotional valence associated with different taste qualities (Pizzagalli et al., 2005; Yang et al., 2023). Therefore, the discriminative capacity observed in the α band may partly reflect its sensitivity to both the sensory and motivational dimensions of gustatory processing.

Beyond the low- and mid-frequency bands, higher-frequency oscillations also contributed meaningfully to taste classification. Although the β and γ bands exhibited slightly lower classification accuracies of 89.22% and 88.89% respectively, their contributions remain notable. The β band is commonly associated with sustained attention, motor preparation, and decision-making processes (Zhang et al., 2008), whereas γ-band activity is associated with fine-grained sensory integration, reward processing, and higher-order cognitive evaluations (HajiHosseini et al., 2012; Yang et al., 2025). In the context of taste perception, β and γ rhythms may potentially reflect the motivational engagement and cognitive appraisal stages following primary sensory encoding (HajiHosseini et al., 2012; Marco-Pallares et al., 2008).

ROC curves in Fig. 6 further corroborate these findings. Consistent with the classification metrics reported in Table 2, the macro-average area under the curve (AUC) values demonstrate that δ and θ bands achieved relatively higher separability between taste categories, with macro-average AUC of 0.94 and 0.93, respectively. The α band followed closely with a macro-average AUC of 0.92, further supporting its important role in gustatory discrimination. Although the β and γ bands exhibited slightly lower macro-average AUC (0.91), their ROC curves still showed robust classification capability. Importantly, the combined multi-frequency feature set achieved the best overall performance, yielding a macro-average AUC of 0.95, consistent with the highest observed accuracy (94.33%) and F1-score (91.37%).

**Fig. 6.**
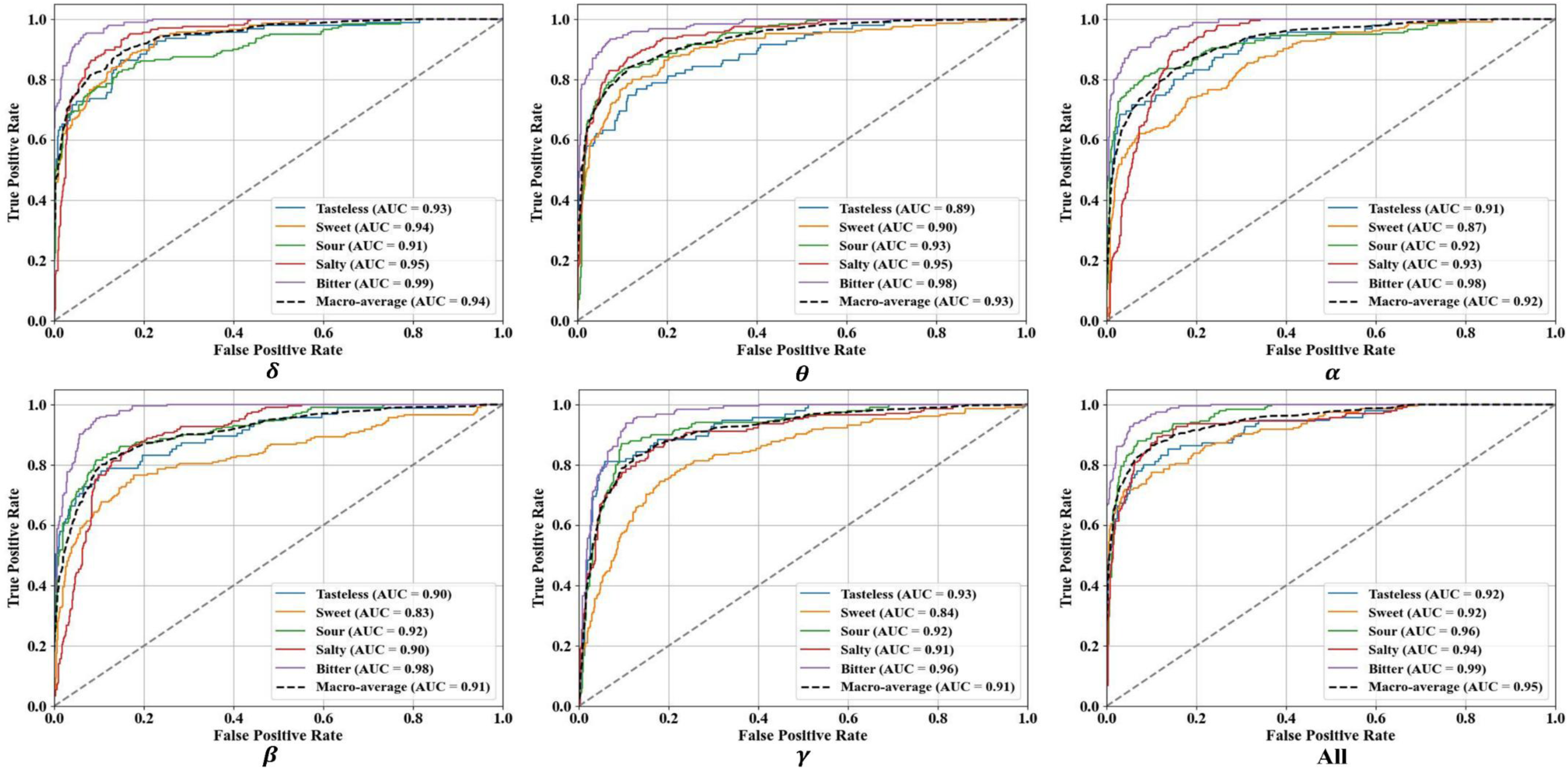
ROC curves for single-frequency and multi-frequency EEG features across different taste categories. AUC values quantify the model’s ability to distinguish between different taste classes, with higher AUC indicating better classification performance.

Taken together, these results indicate the hierarchical neural representations underlying taste perception (Rolls, 2019; Shepherd, 2011). Low-frequency δ and θ rhythms are thought to primarily encode early-stage sensory detection and attentional orientation, mid-frequency α rhythms may integrate sensory inputs with emotional valence appraisal, and higher-frequency β and γ rhythms appear to support evaluative, motivational, and action-related processes. This hierarchical spectral architecture underscores the complex, multi-dimensional nature of gustatory neural processing (Beauchamp, 2019). Our findings are consistent with and extend prior taste-EEG research. Wallroth et al. (2018) demonstrated delta encoding of taste information, aligning with our superior δ-band performance (92.44% accuracy). Similarly, taste-induced modulation of α and θ rhythms reported by Yang et al. (2023) and Kotini et al. (2016) corroborates our α-band and θ-band findings (91.11% and 92.33% accuracy, respectively). Critically, our systematic multi-band analysis reveals the relative contributions across the full spectrum, demonstrating that integrated modeling (94.33% accuracy) surpasses single-band approaches. By integrating features across the full spectral range, the proposed MHASNet model effectively captures the distributed and multifaceted neural signatures underlying taste perception, establishing a comprehensive neurophysiological framework for decoding human gustatory experience from EEG signals.

#### 3.3.2 Unveiling region-wise cortical contributions of taste perception via MHASNet

To further elucidate the neural mechanisms underlying taste perception and validate the spatial discriminability captured by MHASNet, we conducted a region-wise classification analysis across five cortical areas: the frontal, central, temporal, parietal, and occipital lobes. As shown in Fig. 7a, classification accuracy, F1-score, precision, and recall varied significantly across these regions, indicating functional specificity of taste-evoked EEG signals in the spatial domain.

**Fig. 7.**
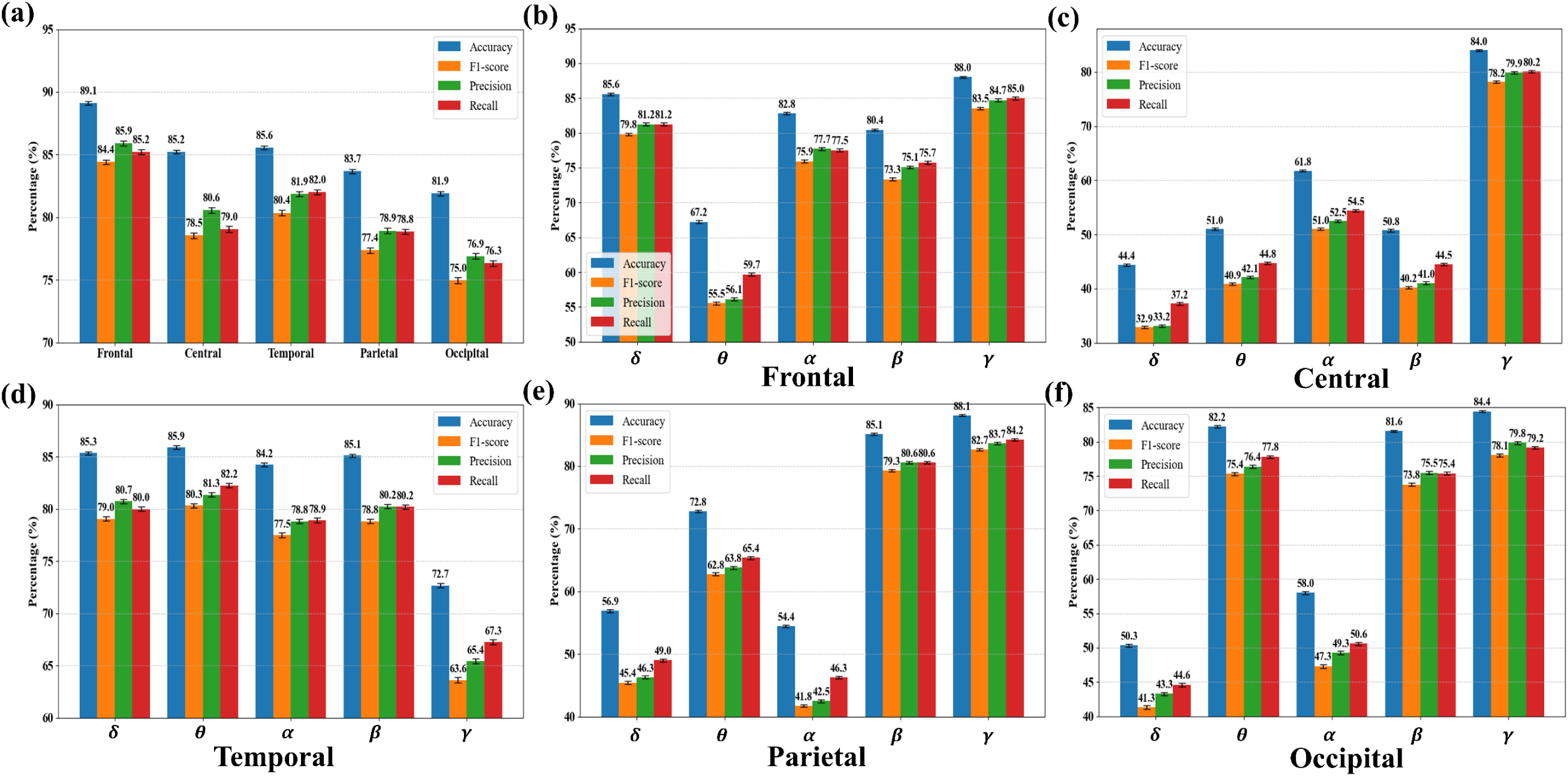
Classification results for different brain regions and different frequency bands. (a) Overall classification results by cortical region: frontal, central, temporal, parietal, and occipital. (b–f) Classification performance for individual frequency bands (δ, θ, α, β, γ) within each brain region.

The frontal region exhibited the most prominent performance, achieving an accuracy of 89.1% and an F1-score of 84.4% (Fig. 7a and Fig. S3a). Its frequency response pattern (Fig. 7b and Fig. 8a) showed strong discriminative power not only in the low-frequency δ band but also in the mid-to-high frequency α, β, and γ bands. This multi-band enhancement is consistent with the established role of the frontal cortex, particularly the prefrontal and orbitofrontal areas, in gustatory integration, reward evaluation, and emotional regulation (Rolls, 2020; Yang et al., 2023, 2025). Previous studies have shown that mid-to-high frequency rhythms in the frontal lobe are involved in the modulation of pleasantness, value judgment, and attentional allocation, suggesting that the frontal cortex plays a key role in the rapid encoding of affective valence (e.g., sweet: pleasant or bitter: aversive) triggered by taste stimuli (Kringelbach, 2005; Rolls, 2012; Yang et al., 2023).

**Fig. 8.**
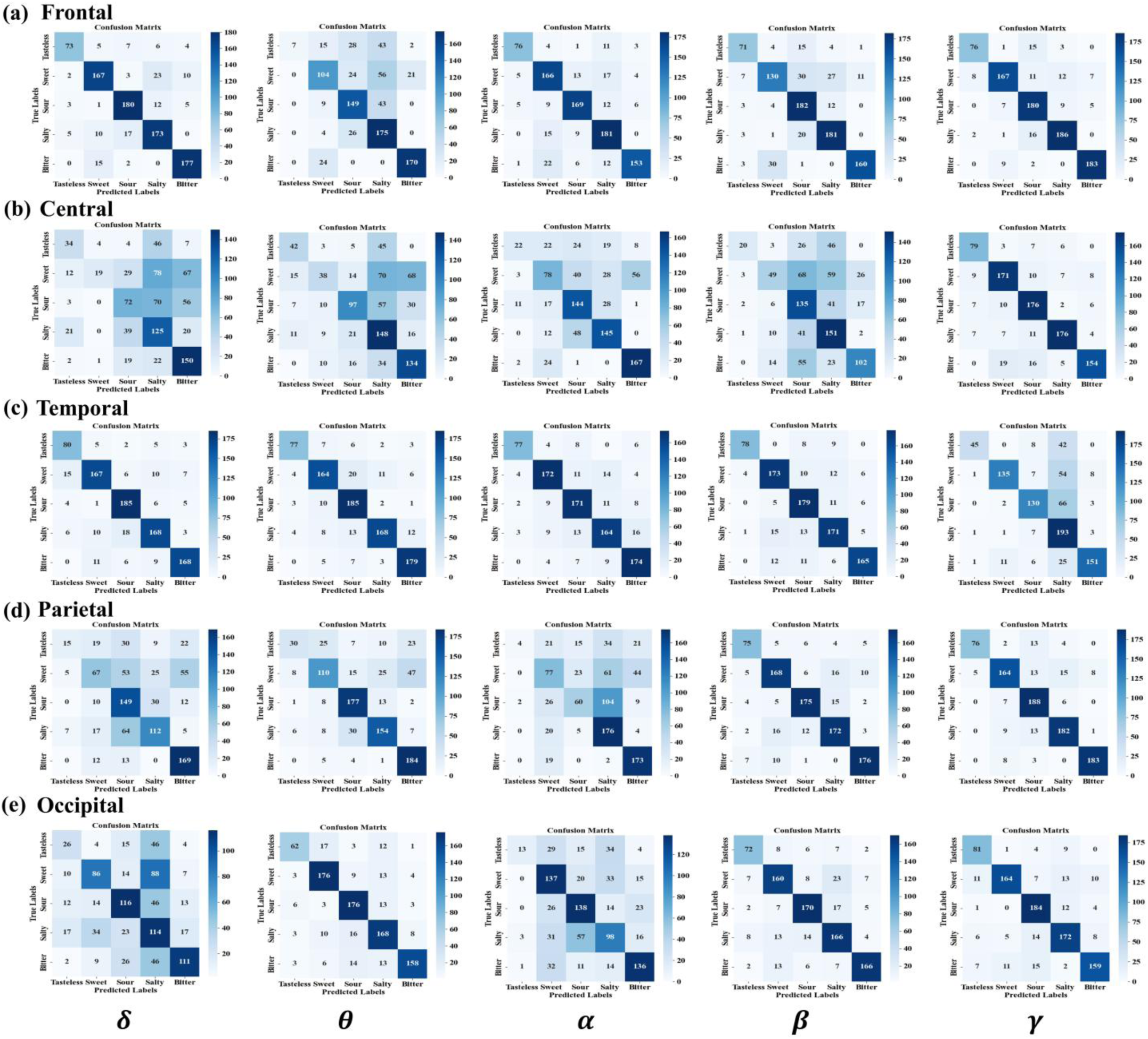
Confusion matrices illustrating taste-type discrimination for individual EEG bands within five cortical regions. Panels (a)–(e) correspond to the frontal, central, temporal, parietal, and occipital areas, respectively.

The temporal and central regions also demonstrated stable classification performance, with accuracies of 85.22% and 85.56%, respectively (Fig. 7a and Fig. S3b-S3c). The central region exhibited a dominant high frequency γ response, while lower frequency bands contributed less (Fig. 7c and Fig. 8b). This spectral preference may reflect the central cortex’s role in taste-evoked somatosensory feedback and attentional regulation (Vincis & Fontanini, 2016). As a primary somatosensory hub for oral and lingual input, the central region appears to be highly sensitive to the tactile differences elicited by different taste types and may enable rapid differentiation among various taste categories through high-frequency neural activity (Shepherd, 2011). In contrast, the temporal region presented an opposite spectral pattern, showing strong classification performance in all frequency bands except γ (Fig. 7d and Fig. 8c). This “low-to-mid frequency dominant” profile suggests that, due to its proximity to the primary gustatory cortex, the temporal lobe is more engaged in early sensory detection, feature encoding, and multimodal integration involving memory and emotion, rather than higher-order cognitive evaluation typically associated with γ rhythms (Patel et al., 2023; Smith, 2019).

The parietal and occipital lobes demonstrated relatively lower overall classification performance, with accuracies of 83.67% and 81.89%, respectively (Fig. 7a and Fig. S3d-S3e). However, they still maintained stable discriminative power in the β, and γ frequency bands (Fig. 7e-7f and Fig. 8d-8e), highlighting their consistent contributions across multiple brain regions (Stein & Stanford, 2008). This suggests that although these regions are not primary gustatory areas, they may still reflect taste-evoked neural activity differences through high-frequency oscillations, particularly in processes related to somatosensory feedback, attentional modulation, or cross-modal spatial information integration (Shepherd, 2011; Vincis & Fontanini, 2016).

In summary, these findings indicate that taste perception engages spatially distinct neural mechanisms with clear functional differentiation across cortical regions (Rolls, 2020; Shepherd, 2011). The frontal cortex, known for its role in sensory appraisal and cognitive regulation, contributed significantly across both low- and high-frequency bands, reinforcing its central role in orchestrating complex gustatory processing. The central region showed a high-frequency dominant profile potentially indicative of sensorimotor integration, likely reflecting rapid responses to taste-induced somatosensory input. In contrast, the temporal lobe was primarily associated with low-to mid-frequency activity, supporting its role in early-stage sensory encoding and the integration of taste with memory and affective context. Our region-wise findings complement fMRI studies of taste perception. The prominent frontal lobe contribution (89.1% accuracy) aligns with neuroimaging evidence highlighting orbitofrontal cortex involvement in taste valuation (Kringelbach, 2005; Rolls, 2020). The temporal region’s performance (85.2% accuracy) is consistent with its proximity to primary gustatory cortex in the insula (Patel et al., 2023). The observed spatial-specific patterns suggest that MHASNet-based taste-EEG evaluation model effectively mirrors the neural architecture involved in gustatory perception.

## 4. Limitations and Outlook

The proposed EEG- and deep-learning-based taste evaluation model demonstrates promising accuracy and translational potential. However, several limitations should be acknowledged. First, owing to the absence of large, publicly available databases of taste-evoked neural responses, this study relied on a self-collected dataset with a relatively small sample size and demographically homogeneous participants. While this dataset was sufficient to demonstrate the feasibility of the proposed approach, future validation with larger and more diverse populations will be valuable for enhancing its generalizability across different cultural and dietary backgrounds. Second, the experiments were conducted under controlled laboratory conditions with discrete taste stimuli, which may not fully reflect the complexity of real-world flavor experiences. In addition, the current model primarily focuses on basic taste categories; future research could extend its scope to include taste mixtures and substitutes exhibiting primary and secondary taste qualities (e.g., salt replacers combining saltiness with bitterness, or sweeteners with varying secondary taste profiles), complex flavor profiles, hedonic intensity, and dynamic temporal changes during eating. Addressing these issues in future work will help improve the robustness, ecological validity, and application potential of the proposed method.

Despite these limitations, the systematic analyses presented in this study, revealing hierarchical spectral dynamics and region-specific contributions to taste perception, open several avenues for advancing taste research. The hierarchical organization identified across frequency bands and cortical regions provides a framework for investigating how neural representations evolve with dietary interventions, potentially revealing mechanisms underlying taste plasticity and preference formation. Extending this work through integration with complementary neuroimaging modalities could enable more comprehensive characterization of gustatory processing networks, while exploring connectivity patterns may elucidate how distributed frequency bands and brain regions coordinate to encode complex flavor experiences.

From an applied perspective, the model offers an objective, neurophysiology-based framework that complements subjective sensory evaluations and instrumental analyses for taste evaluation. This approach offers unique advantages including elimination of cultural and linguistic biases inherent in verbal reports, cross-population comparability for international applications, and assessment capabilities for special populations unable to provide reliable verbal feedback. This capability lays the groundwork for taste-oriented brain–computer interfaces and taste-evaluation platforms. Its data-driven approach can inform precision product development, enabling food developers to tailor and optimize flavor attributes based on quantitative neural feedback rather than trial-and-error. By aligning objective neural assessment of taste with industrial workflows, this approach can enhance product-development efficiency, ensure consistent quality, and better meet evolving consumer demands.

## 5. Conclusion

In summary, this study established an EEG-based taste-evaluation model built on MHASNet, offering an objective, neurophysiology-based complement to traditional sensory panel evaluations. We recorded EEG under five taste conditions (sweet, sour, salty, bitter, and tasteless) and assembled a dedicated taste-EEG dataset; the model was then trained to decode taste stimuli from EEG signals. MHASNet integrates multi-band feature extraction, multi-scale temporal convolutions, dual-attention mechanisms, and a squeeze-and-excitation module to adaptively emphasize salient spatiotemporal and spectral features. In comparative and ablation experiments, the model achieved 94.33% accuracy, 91.37% F1-score, 91.89% precision, and 92.07% recall, while maintaining millisecond-level inference latency suitable for real-time applications. Moreover, MHASNet elucidated multidimensional neural responses to gustatory stimulation and provided insight into how different tastes influence consumer sensory responses.

## CRediT authorship contribution statement

**Tianyi Yang:** Conceptualization, Methodology, Formal analysis, Writing-original draft. Writing-review & editing. **Yueyue Xiao:** Data curation, Formal analysis, Investigation, Writing-original draft. **Zhaoyan Li:** Data curation, Formal analysis, Investigation. **Chunxiao Chen:** Resources, Supervision. **Fei Ye:** Formal analysis, Investigation. **Mian Cao:** Formal analysis, Investigation. **Rui Liu:** Project administration, Supervision. **Yingli Zhu:** Investigation, Writing-review & editing. **Zhiyu Qian:** Project administration, Resources, Funding acquisition, Supervision. **Jagath C. Rajapakse:** Resources, Writing-review & editing.

## Acknowledgments

The authors would like to thank all the volunteers who participated in this study.

